## Supplementary Figures for "*Pfcerli2*, a duplicated gene in the malaria parasite *Plasmodium falciparum* essential for invasion of erythrocytes as revealed by phylogenetic and cell biological analysis"

SUPPLEMENTARY INFORMATION

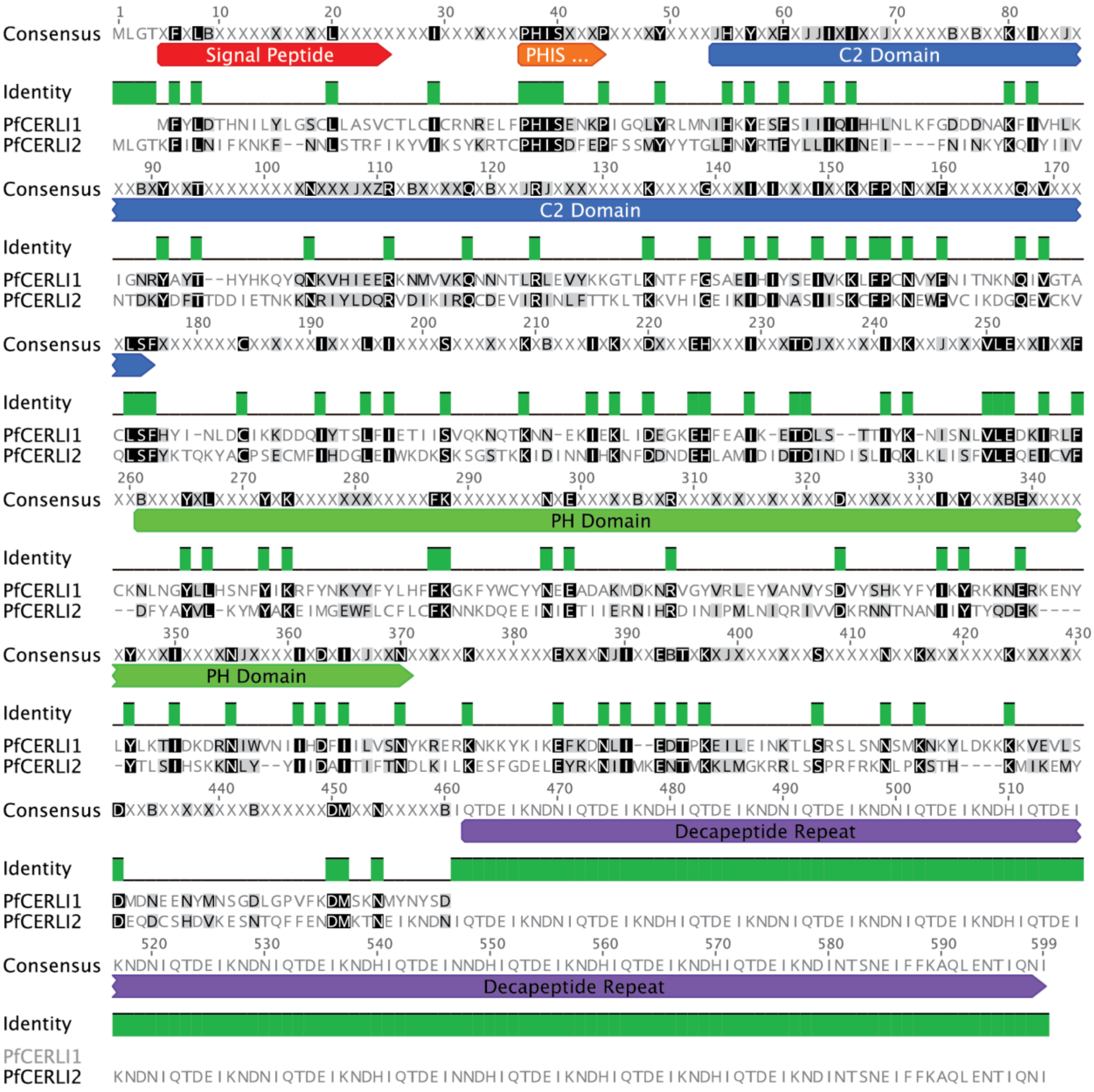


Supplementary Figure 1: Alignment of PfCERLI1 and PfCERLI2.

*Geneious global alignment with free-end gaps between the amino acid sequences of PfCERLI1 (Pf3D7_0210600) and PfCERLI2 (Pf3D7_0405200). The signal peptide is predicted only for PfCERLI1 and is represented here as it is currently annotated on PlasmoDB. Removal of this signal peptide region does not affect localisation of PfCERLI1*[1]*. The PH domain is predicted only for PfCERLI1, and not PfCERLI2.*

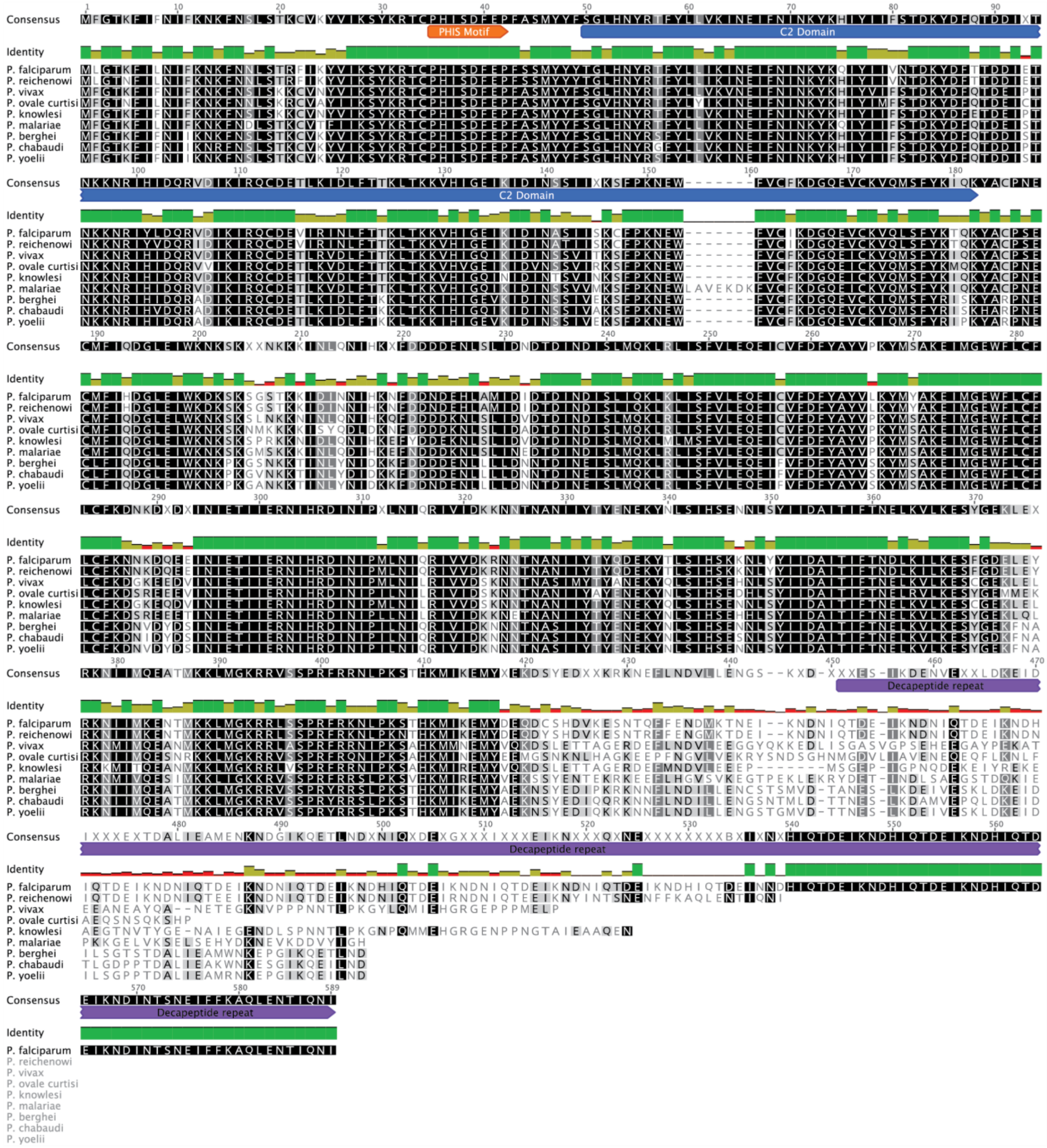


Supplementary Figure 2: Alignment of PfCERLI2 and its *Plasmodium* homologues.

*Geneious global alignment with free-end gaps between the amino acid sequences of PfCERLI2 (Pf3D7_0405200) and its homologues in P. reichenowi (PRCDC_0402800), P. vivax (PVP01_0304600), P. ovale curtisi (PocGH01_030212700), P. knowlesi (PKNH_0303300) , P. malariae (PmUG01_03015300), P. berghei (PBANKA_1002900), P. chabaudi (PCHAS_1003800) , and P. yoelii (PY17X_1004300).*

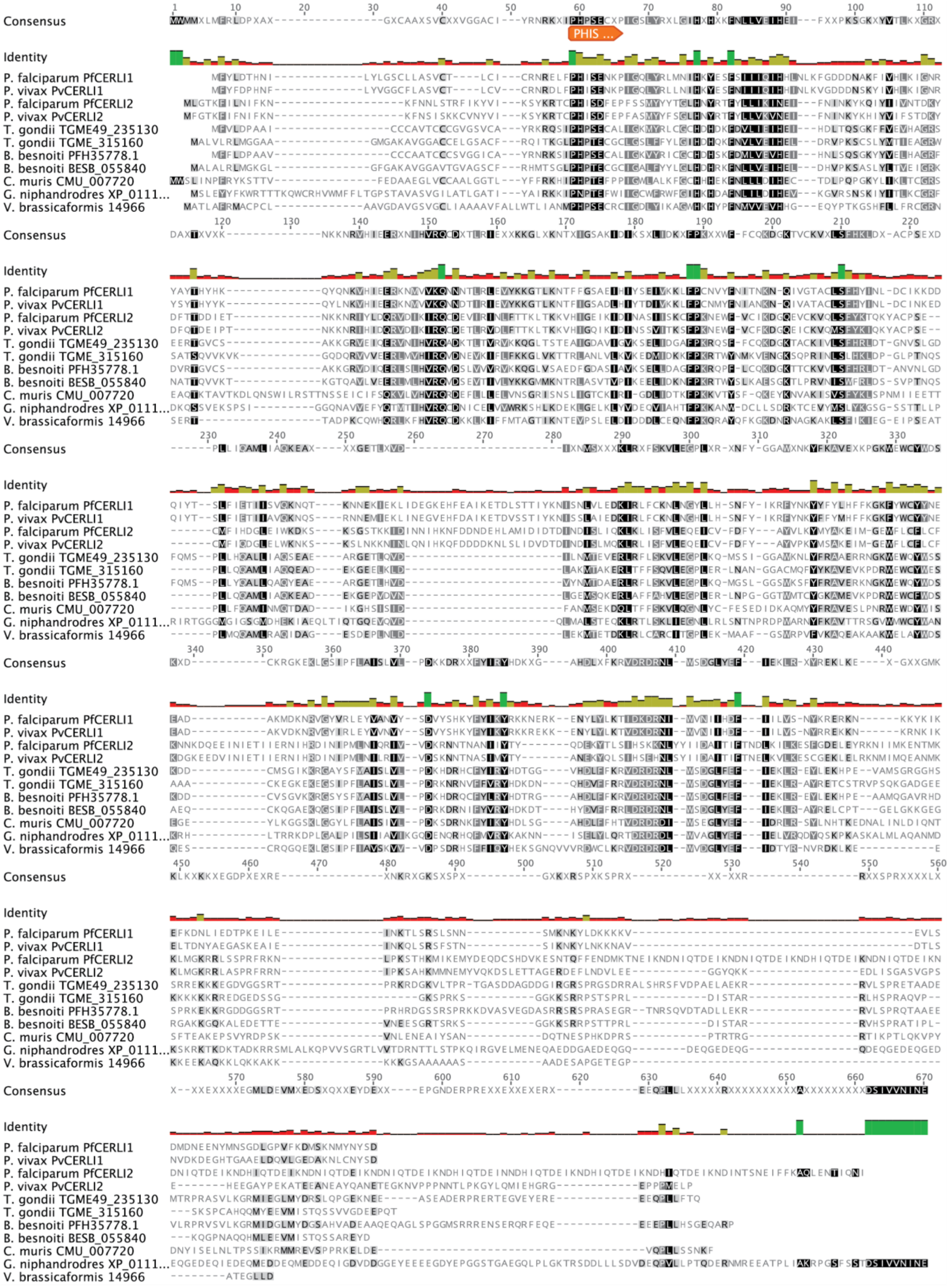

Consensus
PHIS ...
Identity
P. falciparum PfCERLI1
P. vivax PvCERLI1
P. falciparum PfCERLI2
P. vivax PvCERLI2
T. gondii TGME49_235130
T. gondii TGME_315160
B. besnoiti PFH35778.1
B. besnoiti BESB_055840
C. muris CMU_007720
G. niphandrodes XP_0111...
V. brassicaformis 14966

Supplementary Figure 3: Alignment of PfCERLI1, PfCERLI2, and their Apicomplexan homologues.

*Geneious global alignment with free-end gaps between the amino acid sequences of PfCERLI1 (Pf3D7_0210600), PfCERLI2 (Pf3D7_0405200) and their homologues in P. vivax (PVP01_0414300 & PVP01_0304600), Toxoplasma gondii (TGME49_235130 & TGME49_315160), Besnoitia besnoiti (PFH35778.1 & BESB_055840), Cryptosporidium muris (CMU_007720) , Gregarina niphandrodes (XP_011129839.1), and Vitrella brassicaformis (14966).*

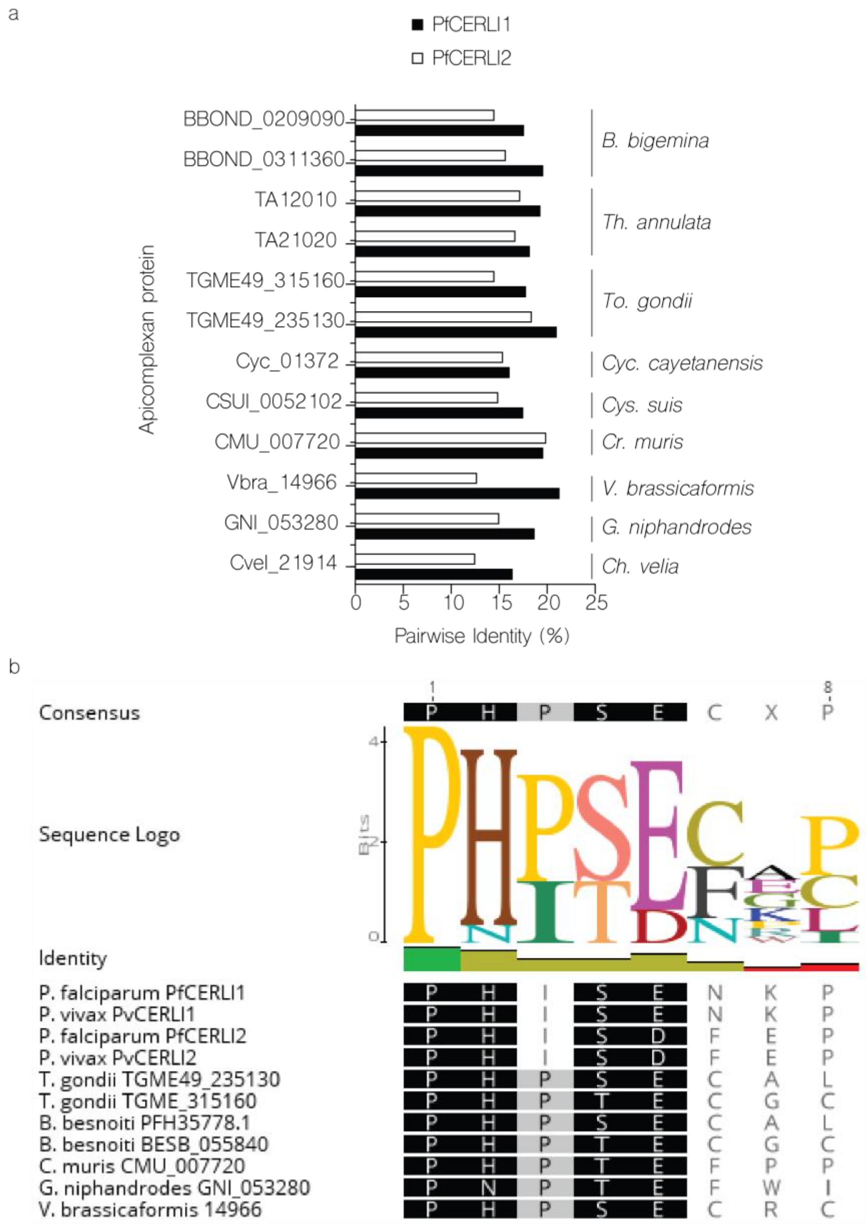


Supplementary Figure 4: Pairwise identity of PfCERLI1, PfCERLI2, and their Apicomplexan homologues.

***(a)*** *Pairwise identities, as determined by Geneious global alignment with free-end gaps, of the amino acid sequence for either PfCERLI1 or PfCERLI2 and their homologues in Babesia bigemina (BBOND_0209090 & BBOND_0311360), Theileria annulata (TA12010 & TA21020), Toxoplasma gondii (TGME49_235130 & TGME49_315160), Cyclospora cayetanensis (Cyc_01372), Cystoisospora suis (CSUI_0052102), Cryptosporidium muris (CMU_007720), Vitrella brassicaformis (Vbra_14966), Gregarina niphandrodes (GNI_053280), and Chromera velia (Cvel_21914).* ***(b)*** *Consensus sequence and conservation of the PHIS motif in PfCERLI1, PfCERLI2, and their apicomplexan homologues.*

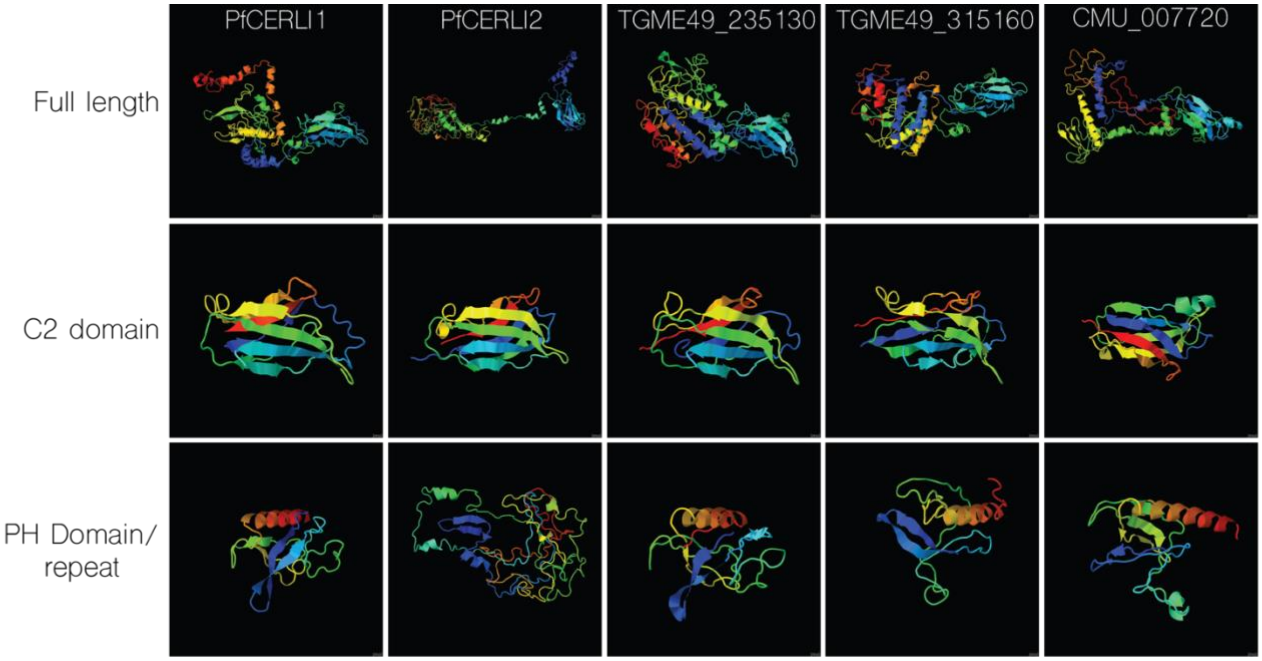


Supplementary Figure 5: Predicted protein structures of CERLI1, CERLI2 and their homologues.

*Full length protein structure of PfCERLI1, PfCERLI2, their T. gondii homologues (TGME49_235130 and TGME49_315160), and their Cryptosporidium muris homologue (CMU_007720) was predicted using Phyre2. All proteins were predicted to have a C2 domain, and all proteins except PfCERLI2 were also predicted to contain a Pleckstrin homology (PH) domain.*

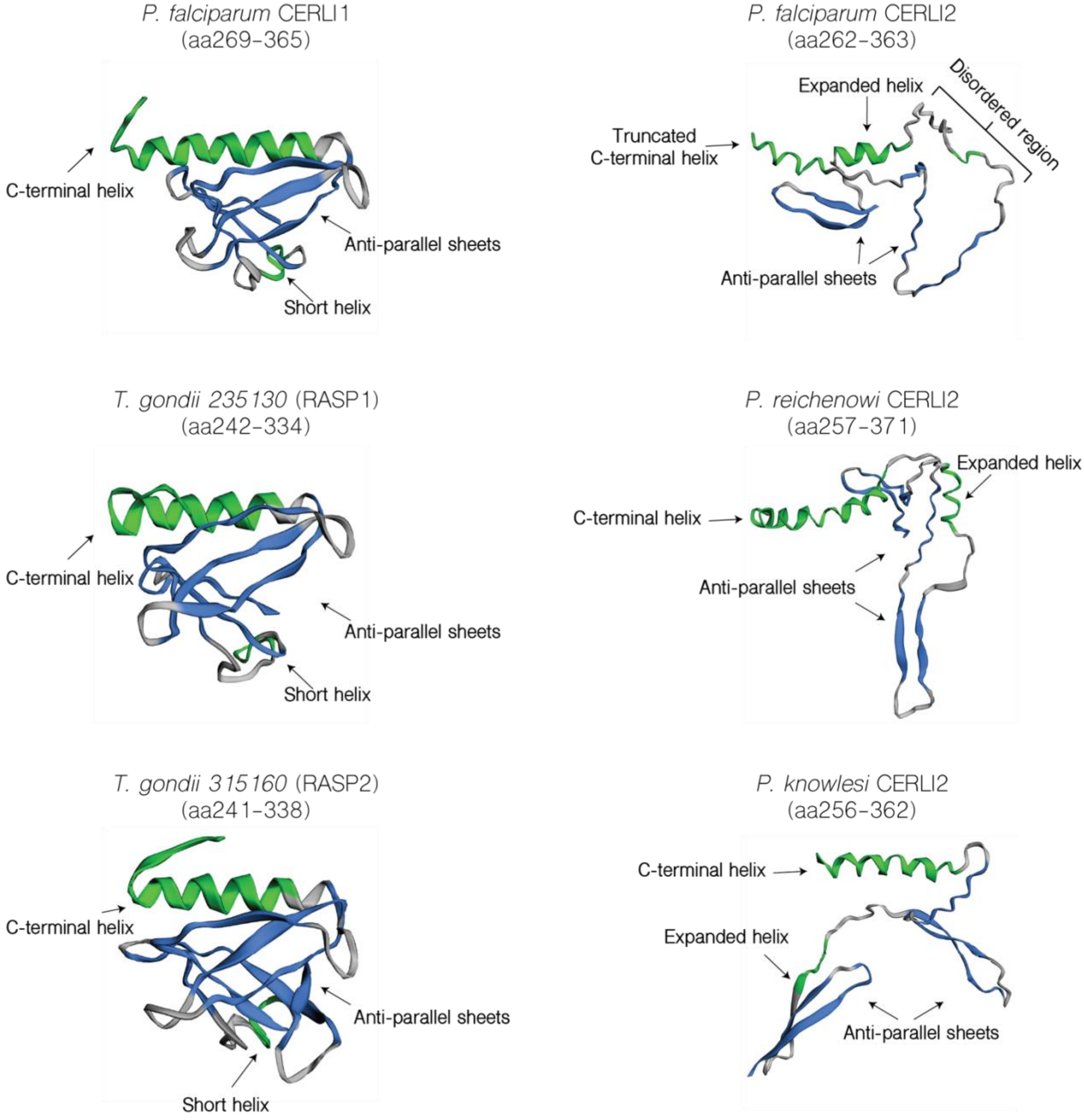


Supplementary Figure 6: Predicted structure of the PH domain region of PfCERLI2, PfCERLI1 and their homologues.

*Full length protein sequences for PfCERLI1, PfCERLI2, along with both their homologues in Toxoplasma gondii, and the homologues of CERLI2 in Plasmodium reichenowi and Plasmodium knowlesi, were predicted using Phyre2. The PH domain region of those structures was visualised using EzMol. Region of each protein depicted are shown in brackets. Green = alpha-helix, blue = beta-strand, grey = coil.*

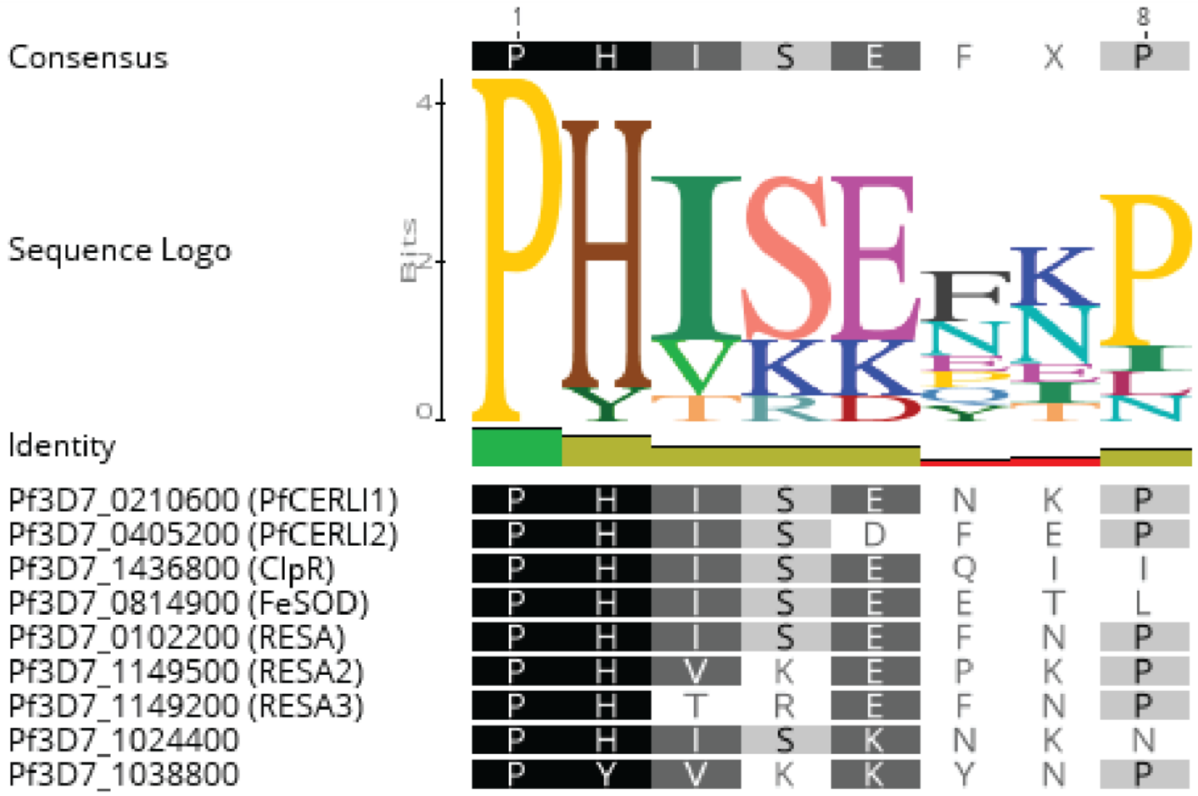


Supplementary Figure 7: PHIS consensus sequence of *P. falciparum* PHIS-containing proteins.

*Consensus sequence and conservation of the PHIS motif for PHIS-containing proteins of P. falciparum.*

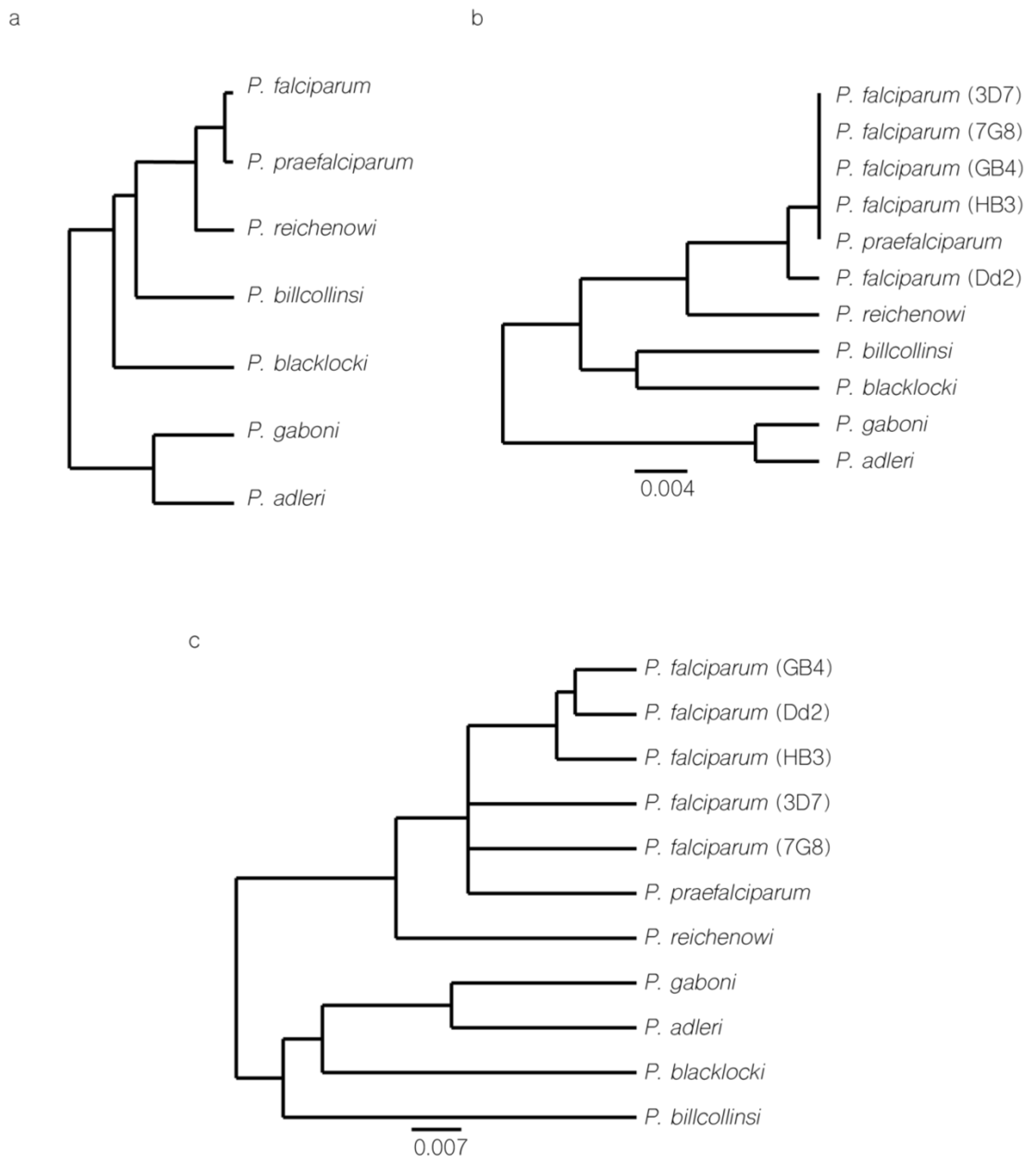


Supplementary Figure 8: The decapeptide repeat of PfCERLI2 is under differential selection in *Laverania*.

***(a)** Cladogram of relationships between Laverania as determined by Otto et al., Nat. Microb. 2018. **(b)** Phylogenetic tree of PfCERLI2 and its homologues in Laverania and multiple P. falciparum isolates when the decapeptide repeat region of each protein has been removed. **(c)** Phylogenetic tree of full length PfCERLI2 and its homologues in Laverania and multiple P. falciparum. Scale bars = amino acid substitutions per site. Phylogenetic trees constructed using unweighted pair group method with arithmetic mean (UPGMA). Branch length corresponds to amino acids substitutions per site.*

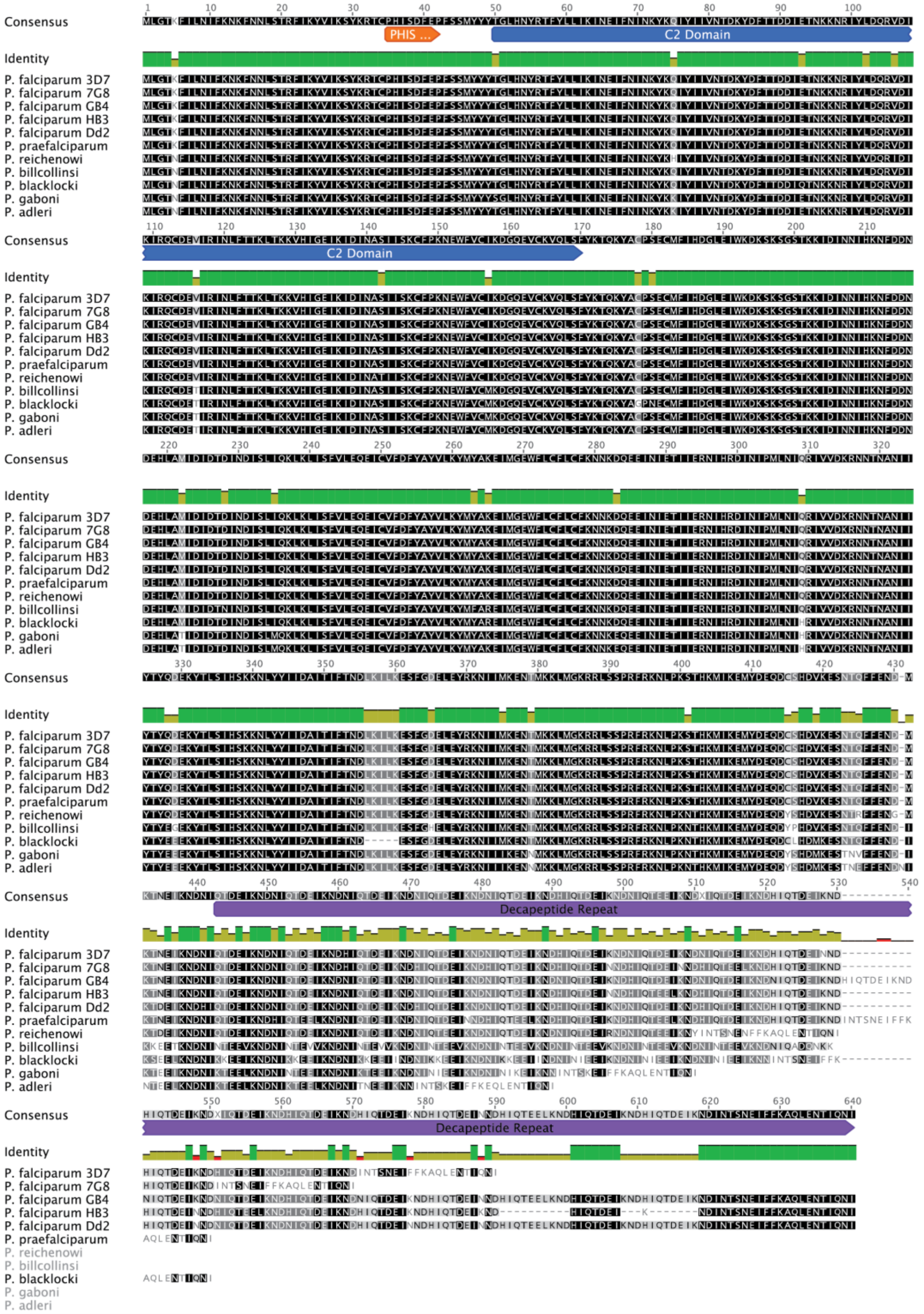


Supplementary Figure 9: Alignment of PfCERLI2 and its homologues in Laverania.

*Geneious global alignment with free-end gaps between full length PfCERLI2 in Laverania and multiple P. falciparum isolates.*

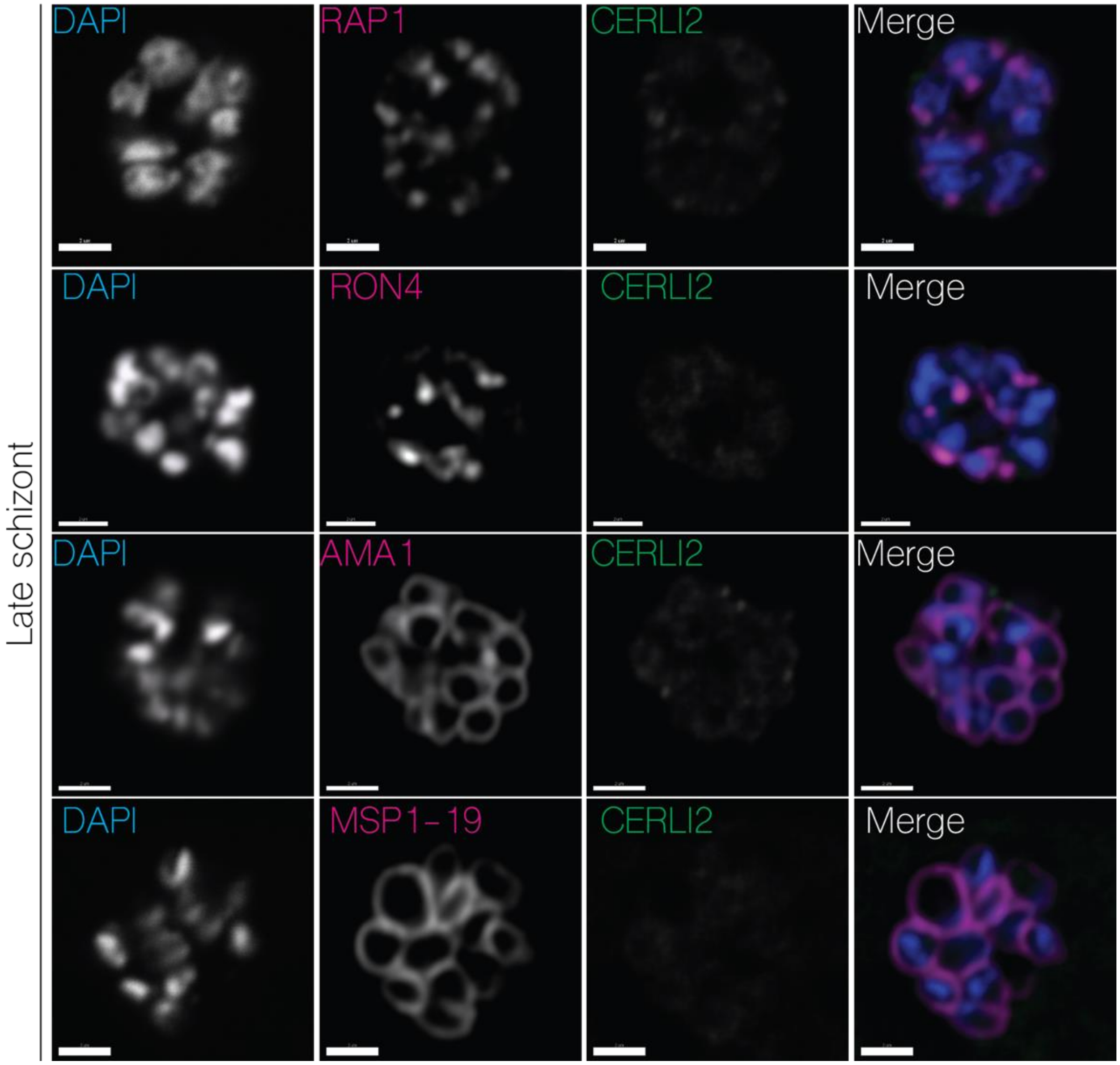


Supplementary Figure 10: Immunofluorescence microscopy of PfCERLI2 in late schizonts shows reduced CERLI2 signal.

*Immunofluorescence microscopy of either late* $PfCERLI2^{HAGlmS}$ *schizonts stained with DAPI (nucleus) and anti-HA (PfCERLI2) antibodies, along with antibodies to either RAP1 (rhoptry bulb, RON4 (rhoptry neck), AMA1 (micronemes), or MSP1-19 (merozoite surface). Scale bar = 2 µm*

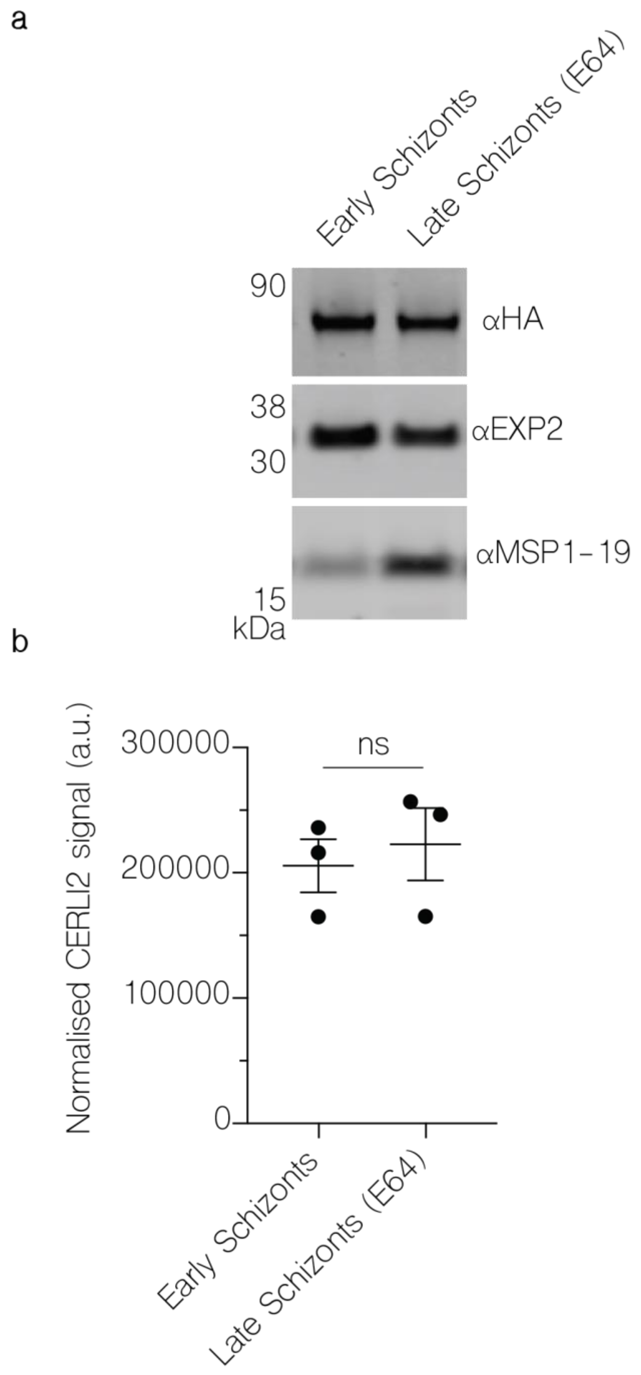


Supplementary Figure 11: Western blot of early and late PfCERLI2$^{HAGlmS}$ schizonts.

***(a)*** *PfCERLI2$^{HAGlmS}$ parasite lysates were either harvested for Western blot as early schizonts (~44 hours post invasion), or matured in the presence of E64 and harvested as late schizonts (~48). Lysates were probed with either anti-HA (PfCERLI2), anti-EXP2 (loading control), or anti-MSP1-19 (schizont maturity control) antibodies. Image representative of 3 biological replicates.* ***(b)*** *Quantification of PfCERLI2 signal, normalised to EXP2, for both early and late schizonts. n=3, ns=p>0.05 by unpaired t-test. Error bars = SEM.*

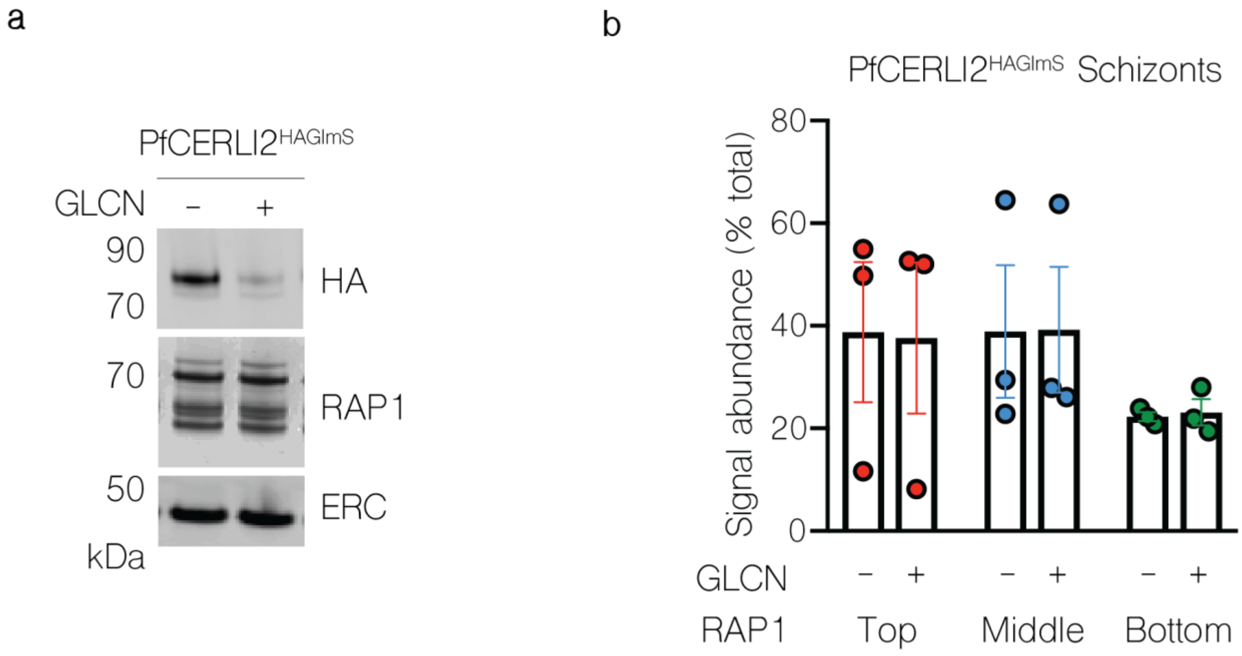


Supplementary Figure 12: PfCERLI2 knockdown does not alter RAP1 processing in C1 arrested schizonts.

***(a)*** *PfCERLI2$^{HAGlmS}$ parasites were either treated with 2.5 mM GLCN or left untreated, before being matured in the presence of C1 to prevent PVM rupture. Parasite lysates were then prepared and probed with anti-HA (PfCERLI2), anti-RAP1 (rhoptry bulb), and anti-ERC (loading control) antibodies. Images representative of 3 biological replicates.* ***(b)*** *Quantification of individiual band intensities, as a percentage of the total signal, from RAP1 signals using the parasite lysates from the secretion assay. n=3 biological replicates. Error bars = SEM.*

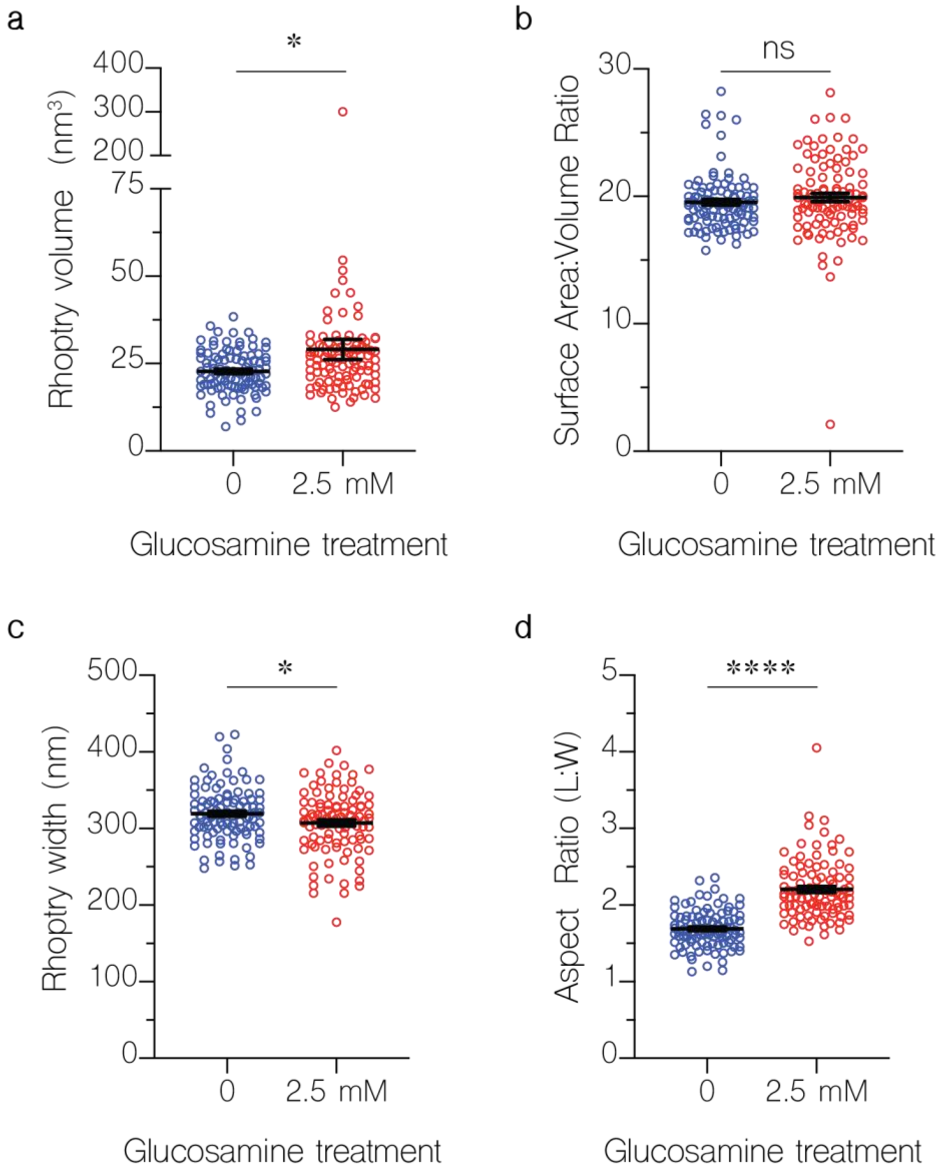


Supplementary Figure 13: PfCERLI2 knockdown grossly alters rhoptry morphology.

*PfCERLI2$^{HAGlmS}$ parasites were either treated with 2.5 mM GLCN, or left untreated, from ring-stages and arrested at schizont stages using C1. These parasites were then imaged by serial block face scanning electron microscopy (SBF-SEM) array tomography. 100 rhoptries for each treatment were segmented, with their volume* ***(a)****, surface are to volume ratio* ***(b)****, width* ***(c)****, and length to width aspect ratio* ***(d)*** *calculated. Error bars = SEM. ns = $p>0.05$, *=$p<0.05$, ****=$p<0.0001$ by unpaired t-test.*

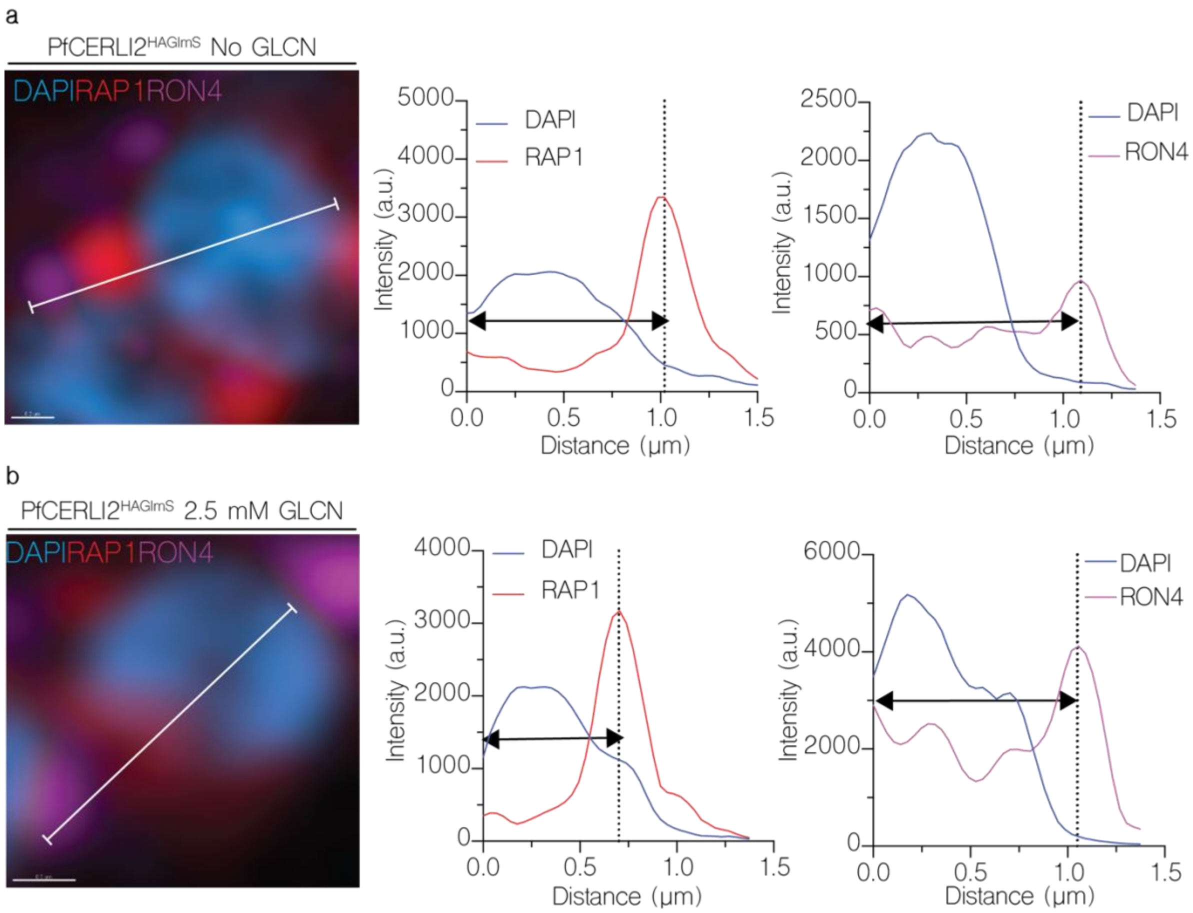


Supplementary Figure 14: Knockdown of PfCERLI2 alters rhoptry antigen positioning and distribution.

*PfCERLI2$^{HAGlmS}$ schizonts were matured in the presence of E64, stained with antibodies against RAP1 (rhoptry bulb) and RON4 (rhoptry neck), and imaged by Airyscan super-resolution microscopy. The flourescence intensity from maximum-intensity projections of RAP1 and RON4 signals were then measured from the basal end of the nucleus and plotted against merozoite length for parasites that were either left untreated* ***(a)*** *or treated with 2.5 mM GLCN* ***(b)****. Dashed lines = RAP1/RON4 maximum fluorescence intensity.*

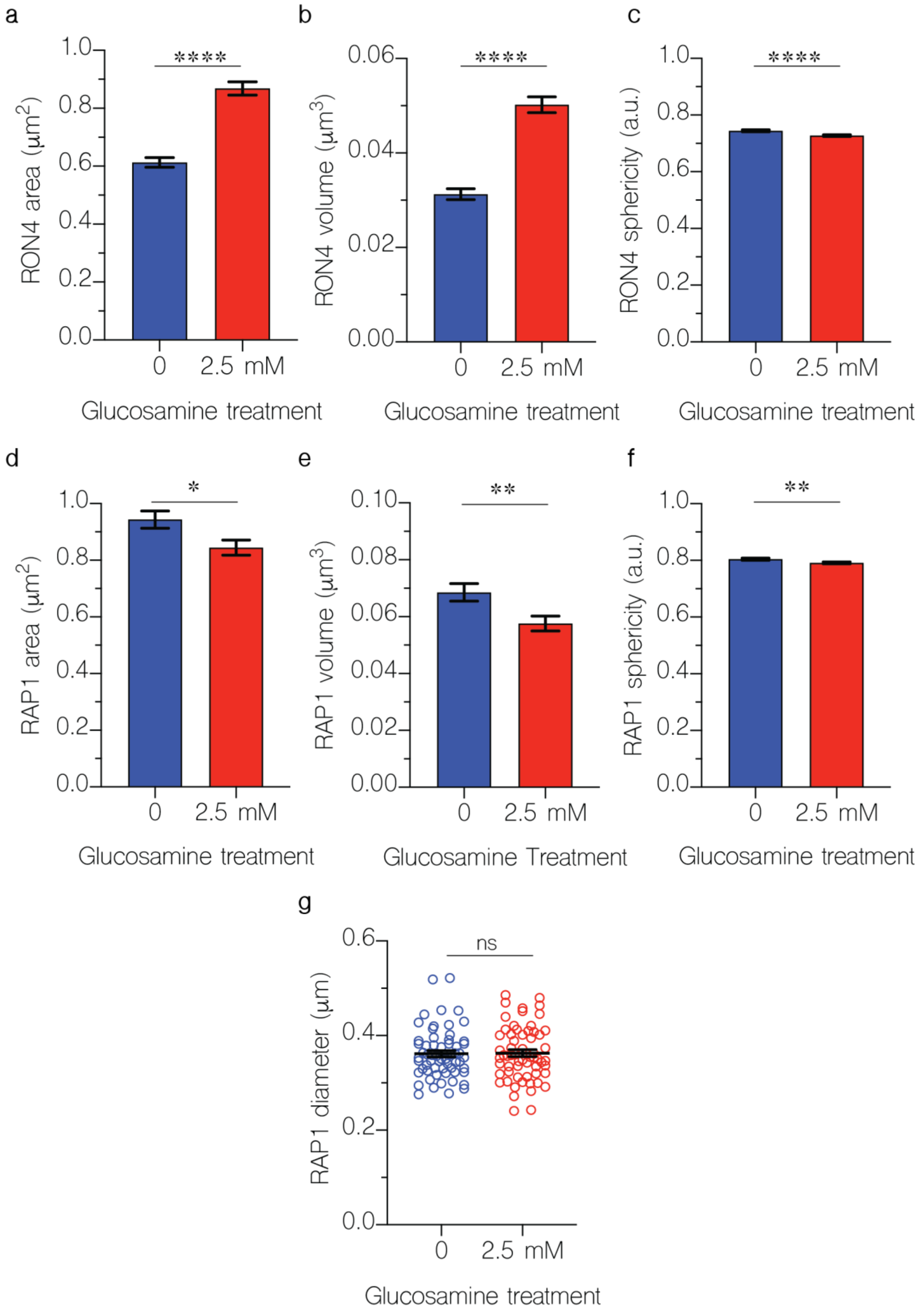
a
RON4 area (μm2)
1.0
0.8
0.6
0.4
0.2
0.0
****
0
2.5 mM
Glucosamine treatment
b
RON4 volume (μm3)
0.06
0.04
0.02
0.00
****
0
2.5 mM
Glucosamine treatment
c
RON4 sphericity (a.u.)
1.0
0.8
0.6
0.4
0.2
0.0
****
0
2.5 mM
Glucosamine treatment
d
RAP1 area (μm2)
1.0
0.8
0.6
0.4
0.2
0.0
*
0
2.5 mM
Glucosamine treatment
e
RAP1 volume (μm3)
0.10
0.08
0.06
0.04
0.02
0.00
**
0
2.5 mM
Glucosamine Treatment
f
RAP1 sphericity (a.u.)
1.0
0.8
0.6
0.4
0.2
0.0
**
0
2.5 mM
Glucosamine treatment
g
RAP1 diameter (μm)
0.6
0.4
0.2
0.0
ns
0
2.5 mM
Glucosamine treatment

Supplementary Figure 15: PfCERLI2 knockdown alters the distribution of rhoptry markers.

*PfCERLI2$^{HAGlmS}$ parasites were either treated with 2.5 mM GLCN from ring-stages until schizont stages, or left untreated. Schizonts were then matured in the presence of the egress inhibitor E64. Parasites were then stained with DAPI, anti-RAP1 (rhoptry bulb), and anti-RON4 (rhoptry neck) antibodies, and analysed by Airyscan super-resolution microscopy. RON4 foci area* ***(a)****, volume* ***(b)*** *and sphericity* ***(c)****, along with RAP1 foci area* ***(d)****, volume* ***(e)****, and sphericity* ***(f)*** *were then quantified using an established automated image analysis pipeline for these markers[1]. 779 foci quantified for RON4 untreated, 1042 foci quantified for RON4 + 2.5 mM GLCN, 367 foci quantified for RAP1 untreated, 386 foci quantified for RAP1 + 2.5 mM GLCN.* ***(g)*** *Images of the same parasites were then blinded and the rhoptry bulb diameter (RAP1 signal) measured, with each datapoint representing a single rhoptry. 60 rhoptries were measured for untreated parasites, and 62 for + 2.5 mM GLCN parasites. N=3, all error bars = SEM, ns = $p>0.05$, * = $p<0.05$, ** = $p<0.01$, **** = $p<0.0001$ by unpaired t-test.*
